# Mcr-dependent methanogenesis in *Archaeoglobaceae* enriched from a terrestrial hot spring

**DOI:** 10.1101/2022.12.01.518715

**Authors:** Steffen Buessecker, Grayson L. Chadwick, Melanie E. Quan, Brian P. Hedlund, Jeremy A. Dodsworth, Anne E. Dekas

## Abstract

The preeminent source of biological methane on Earth is methyl coenzyme M reductase (Mcr)-dependent archaeal methanogenesis. A growing body of evidence suggests a diversity of archaea possess Mcr, however, experimental validation of hypothesized methane metabolisms has been missing. Here, we provide evidence of a functional Mcr-based methanogenesis pathway in a novel member of the family *Archaeoglobaceae*, designated *Methanoproducendum nevadense*, which we enriched from a terrestrial hot spring on the polysaccharide xyloglucan. Our incubation assays demonstrate methane production that is highly sensitive to the Mcr-inhibitor bromoethanesulfonate, stimulated by xyloglucan and xyloglucan-derived sugars, concomitant with the consumption of molecular hydrogen, and causing a deuterium fractionation in methane characteristic of hydrogenotrophic and methylotrophic methanogens. Combined with the recovery and analysis of a high-quality *M. nevadense* metagenome-assembled genome encoding a divergent Mcr and diverse potential electron and carbon transfer pathways, our observations suggest methanogenesis in *M. nevadense* occurs via Mcr and is fueled by the consumption of cross-fed byproducts of xyloglucan fermentation mediated by other community members. Phylogenetic analysis shows close affiliation of the *M. nevadense* Mcr with those from Korarchaeota, Nezhaarchaeota, Verstraetearchaeota, and other *Archaeoglobales* that are divergent from well-characterized Mcrs. We propose these archaea likely also use functional Mcr complexes to generate methane on the basis of our experimental validation in *M. nevadense*. Although our stable isotope approach reveals that microbial methanogenesis contributes only a small proportion of the overall methane abundance in the native habitat, divergent Mcr-encoding archaea may be underestimated sources of biological methane in terrestrial and marine hydrothermal environments.

## Introduction

Biological methane has shaped the past (Pavlov *et al*., 2001; Zahnle, 2006) and present (Arias *et al*., 2021) planetary climate as an agent of atmospheric warming. However, the full diversity and abundance of microorganisms responsible for biological methane production is still not entirely understood. Earth’s major methane source is methyl coenzyme M reductase (Mcr)-dependent metabolism in anaerobic archaea. This 300 kilodalton protein complex catalyzes the terminal step in methanogenesis (Ellefson and Wolfe, 1980; Ermler *et al*., 1997) as well as the initial step in anaerobic methane oxidation (Hallam *et al*., 2004). Divergent homologs of the Mcr complex have been identified in metagenome-assembled genomes (MAGs) of uncultured archaea, including the Bathyarchaeota, Verstraetearchaeota, *Methanonatronarchaeia*, Nezhaarchaeota, *Nitrososphaerales* (syn. Thaumarchaeota), Hadarchaeota, *Archaeoglobales*, ANME-1 and GoM-Arc1 (Evans *et al*., 2015; Borrel *et al*., 2019; McKay *et al*., 2019; Hua *et al*., 2019). The divergence of these new members of the Mcr family appears to be associated with a transition from methane metabolism to short-chain alkanes, such as ethane (Chen *et al*., 2019; Hahn *et al*., 2020) and butane (Laso-Pérez *et al*., 2016), leading them to be referred to as alkyl-CoM reductases (Acrs). The explosion of MAGs from uncultured archaeal lineages containing Mcr-and Acr-encoding sequences in the last few years has revealed a large uncertainty in our understanding of methane and alkane metabolisms.

Recently, Mcr/Acr complexes have been uncovered in MAGs corresponding to members of the *Archaeoglobi*, cultured representatives of which are not known to catalyze methanogenesis or methanotrophy. One such genome from *Candidatus* Polytropus marinifundus, derived from the deep subseafloor along the Juan de Fuca Ridge (Boyd *et al*., 2019), contained two putative Acr operons. This archaeon was hypothesized to use these enzymes for the oxidation of alkanes, similar to other Acr-encoding archaea (Chen *et al*., 2019; Hahn *et al*., 2020; Laso-Pérez *et al*., 2016). Despite several reports of small amounts of methane (< 200 µmol L^−1^ culture) produced by *Archaeoglobus* isolates (Stetter *et al*., 1987; Beeder *et al*., 1994; Huber *et al*., 1997; Mori *et al*., 2008), a role in methane metabolism in this organism seemed unlikely in light of the high-degree sequence divergence from experimentally validated Mcrs and, moreover, methanogenesis has been suggested to have been lost in the *Archaeoglobi* (Bapteste *et al*., 2005). In contrast, a survey of metagenomic data from hot springs and oil reservoirs revealed additional, phylogenetically distinct MAGs of the family *Archaeoglobaceae* containing canonical Mcr operons (Liu *et al*., 2020) and, thus, pointing toward the potential for methane metabolism in the class *Archaeoglobi*. However, experimental evidence of significant methane production or consumption in the *Archaeoglobi* in association with a functional Mcr complex has been lacking and their role in methane cycling in their thermal habitat remained unresolved. While methanogenic activity occurs in marine systems at temperatures of up to 122 °C (Takai *et al*., 2008), *in-situ* substrate abundances may limit methanogenesis in terrestrial systems (Hedlund *et al*., 2015) where it has been observed at ≤ 75 °C. In light of the substrate diversity of divergent Mcrs it may, thus, be conceivable that phyla using divergent Mcrs for methane production showed higher temperature resistance.

We investigated methane metabolism in a member of the *Archaeoglobaceae*, here designated *Methanoproducendum nevadense* XG-1, that was enriched in a mixed, anaerobic culture from Great Boiling Spring (GBS), Nevada, while fed with the polysaccharide xyloglucan (XG). After detecting methane in the headspace of the culture, we hypothesized *M. nevadense* to be responsible for methane formation and assessed effects of a suite of different substrates and inhibitors on methane production. Analysis of the medium-quality XG-1 MAG, the only Mcr-encoding member of the enrichment culture, and a conspecific high-quality MAG designated *Methanoproducendum nevadense* GBS^Ts^ from an *in-situ* anaerobic ammonia fiber expansion (AFEX)-treated corn stover enrichment in GBS suggests an Mcr-dependent pathway with electrons sourced from organic compounds in conjunction with H_2_. Our study demonstrates Mcr-based methane production in a member of the *Archaeoglobaceae*, a group previously unknown to include true methanogens, describes the underlying ecology and physiology, and provides a link to biological methane production in a terrestrial hydrothermal setting.

## Methods

### *In-situ* enrichments and laboratory cultivation

The geochemistry and microbiology of Great Boiling Spring (GBS) near Gerlach, Nevada (Costa *et al*., 2009; Cole *et al*., 2013), and Little Hot Creek (LHC) in California’s Long Valley caldera (Vick *et al*., 2010) have been described previously. *In-situ* enrichments on 1 g of ammonia-fiber explosion (AFEX) corn stover sealed in 100 µm nylon mesh bags were performed in the GBS main pool from 26 October 2013 to 28 March 2014 as described in (Peacock *et al*., 2013), which enriched for *Archaeoglobi* before. At LHC, *in-situ* enrichments were deployed the same way from 19 Oct 2013 to 6 April 2014. Transfer to GBS mineral salts medium followed an established approach (Buessecker *et al*., 2022) with the difference that 0.02 % (m/v) xyloglucan served as a sole carbon source and a trace metal solution was added according to Hanada et al. (Hanada *et al*., 1995).

### Genomic DNA extraction

DNA was extracted from the XG-degrading culture by bead-beating with the Fast DNA SPIN Kit for Soil (MP Biomedicals) according to a slightly modified protocol (Buessecker *et al*., 2022).

### 16S rRNA gene tag sequencing

16S rRNA gene tag amplification and sequencing on initial enrichment cultures was performed essentially as described (Kozich *et al*., 2013). A modified forward primer 515 (5’ GTGYCAGCMGCCGCGGTAA) with a Y instead of a C at the 4^th^ position from the 5’ end was used to increase coverage of archaea, and a corresponding change was made to the SeqRead1 primer (Buessecker *et al*., 2022). Sequencing (2×250 bp) was performed on an Illumina iSeq sequencer using the V2 reagent kit and reads were processed with the DADA2 pipeline using the Silva database for taxonomy (Callahan *et al*., 2016).

### Metagenomic sequencing, assembly, analysis and binning of MAGs

Extracted culture DNA was sequenced using the Illumina MiSeq platform (short reads), along with Oxford Nanopore sequencing (long reads). Shotgun metagenome sequencing with Illumina MiSeq (2×250 bp) was performed at Argonne National Laboratories, with library preparation using the Nextera Flex DNA kit. For Oxford Nanopore sequencing, libraries were prepared following the Native Barcoding (EXP-NBD103) and Ligation Sequencing (SQK-LSK108) Kit 1D protocols according to the manufacturer’s instructions, and sequencing was performed on a MinION FLO-MIN106 flow cell (Oxford Nanopore, Oxford Science Park, UK). Reads were mapped to known *Archaeoglobi* reference genomes using BowTie 2 (Langmead and Salzberg, 2012). The mapped short and long reads were assembled with hybridSPAdes (Antipov *et al*., 2016) and the resulting contigs were filtered, merged, and dereplicated. This assembly was then subjected to binning with MetaBAT2 v. 1.7 (Kang *et al*., 2019), as implemented in KBase (Arkin *et al*., 2018). Bins were refined to reduce redundancy and contamination with Anvi’o (Eren *et al*., 2021) and MAG quality and classification was determined using CheckM v1.4.0 (Parks *et al*., 2015) and the GTDB Toolkit v1.1.0 (Chaumeil *et al*., 2020) in KBase. Because the *Archaeoglobaceae* MAG obtained from the enrichment culture was only of medium quality, we compared it to MAGs previously derived from GBS metagenomes by our group using the JSpeciesWS online server (Richter *et al*., 2016). Bin 20 of metagenome 3300005298 in the JGI database was shown to belong to the same species, with an average nucleotide identity (ANI) of 98.4 % and was therefore used for comparative genomics. To screen metagenomic raw reads for *mcrA*, GraftM version 0.13.1 (Boyd *et al*., 2018) was trained using the 499 *mcrA* sequences obtained from NCBI’s PSI-Blast (default configuration) with an e-value of 10^−9^. *McrA* sequences were aligned with IQ-TREE (Trifinopoulos *et al*., 2016) using 0.5 perturbation strength and 1,000 iterations, and the alignment was visualized in the iTOL interface version 6 (Letunic and Bork, 2021).All 76 genomes from the GTDB release 207 in the class *Archaeoglobi* were retrieved and supplemented with four additional genomes from IMG and our incomplete MAG from the xyloglucan enrichment for 80 total genomes. All genomes were assessed for completeness and contamination using CheckM v1.4.0 implemented on the KBase platform. The concatenated, masked alignment of 43 marker genes produced by CheckM was used to generate a phylogenomic tree using RAxML v8.2.12 implemented on XSEDE via the CYPRES Science Gateway v3.3, with the following parameters: protein substitution model: PROTGAMMA, protein substitution matrix: WAG, with 100 iterations of rapid bootstrapping.

### Batch incubation experiments

At the beginning of stationary phase, 100 µL of growing culture was transferred to fresh anoxic media amended with filter-sterilized, anoxic stock solutions added with a flushed syringe. All substrate stocks were prepared using molecular grade reagents and the final concentrations were the following: 0.02 % (w/v) xyloglucan, 0.05 % (w/v) of each monosaccharide (L-fucose, D-galactose, D-xylose), 500 µM of each organic acid (acetate, pyruvate, malate, dimethylsulfoniopropionate), 1 mM 2-methoxybenzoate, and 500 µM trimethylamine. Xyloglucan was added as powder directly to the media. The H_2_/CO_2_ (20:80) mix was added to reach 0.7 bar overpressure. The antibiotic treatment consisted of 100 µg mL^−1^ streptomycin and 100 µg mL^−1^ carbenicillin. The inhibitor bromoethanesulfonate (BES) was added to a final concentration of 10 mM and coenzyme M was added at 140 mg L^−1^. The base medium contained 3.1 mM sulfate and traces of sulfate in a trace metal solution, which was all left out in the –SO_4_^2−^ treatment.

### Methane and hydrogen gas measurements

Culture headspace was sampled with flushed syringes into 12 mL exetainers by replacement with N_2_. Using a xyzTek Bandolero auto sampler, gas from 12 mL exetainers was drawn into a gas chromatograph (Shimadzu GC-2014) equipped with a flame-ionization detector (FID) kept at 250 °C. A 1 m HayeSep N pre-column was serially connected to a 5 m HayeSep-D column conditioned at 75 °C oven temperature. Nitrogen gas (UHP grade 99.999 %, Praxair Inc.) served as carrier with 21 mL min^−1^ flow rate. Methane concentration measurements were calibrated with customized standard mixtures (Scott Specialty Gases, accuracy ± 5 %). Gas phase concentrations were corrected to account for dilution using Henry’s law with the dimensionless concentration constant K^H^_cc_(CH_4_) = 0.0249 for gas dispersion into the aqueous phase at 73 °C.

H_2_ gas concentration in 12 mL exetainer samples was determined with a reactive mercury bed reduced gas analyzer (Peak Laboratories). Gas samples (250 µL) were injected with a gas-tight syringe (VICI) into an N_2_ carrier gas stream through a Unibeads 1S and a Molecular Sieve 13X column conditioned at 104 °C. The retention time for H_2_ was 51 seconds. Blank controls, consisting of uninoculated media with xyloglucan, never produced H_2_ above the detection limit. The analyzer was calibrated using a 100 ppm H_2_ standard (Scott Specialty Gases, accuracy ± 5 %) that was serially diluted with nitrogen gas.

### *In-situ* incubations and gas sampling

To determine methane production rates in the source pool of GBS, as well as nearby source pools G04b and SSW, hot spring sediment was transferred into a glass flask that was continuously flushed with N_2_. Fresh hot spring water was N_2_-sparged and used to dilute the sediment 1:4. The sediment slurry was distributed into pre-flushed vials with a 18G needle. *In-situ* incubations did not receive substrates and the closed, anoxic vials were placed into the spring. Methane production rates were calculated based on the accumulation of methane derived from headspace sampled at the spring. Gas for stable isotope analysis was extracted from water collected with a 60 mL syringe at ∼15 cm depth. Half the syringe was filled with sample water after which the other half was filled with N_2_. The dissolved gas was equilibrated with the N_2_ by shaking the syringe for 10 minutes. The gas phase was then injected into evacuated 12 mL exetainers.

### Stable isotope analysis

Isotopes of C and H in methane gas were analyzed using a Thermo Scientific GasBench-Precon concentration unit interfaced to a Thermo Scientific Delta V Plus isotope ratio mass spectrometer (IRMS) at the UC Davis Stable Isotope Facility and data collection followed the method by Yarnes (Yarnes, 2013).

## Results

### Methane formation in the xyloglucan-degrading culture

After screening diverse thermophilic enrichments for methanogenic activity, we detected methane in the headspace of an anaerobic culture derived from Great Boiling Spring (GBS), Nevada. Another enrichment maintained under identical conditions derived from Little Hot Creek (LHC), California, did not show detectable methane. Both cultures were supplied with XG. The cultures contained comparable abundances of *Candidatus* Caldatribacterium species (11.1-13.9 %) and *Pseudothermotoga* (6.4-12.3 %) but differed particularly in one member belonging to the family *Archaeoglobaceae*, here introduced as *Methanoproducendum nevadense*. Although at a low proportion (0.1 %), *M. nevadense* was present in GBS but not in LHC (Fig. 1A).

**Fig. 1.**
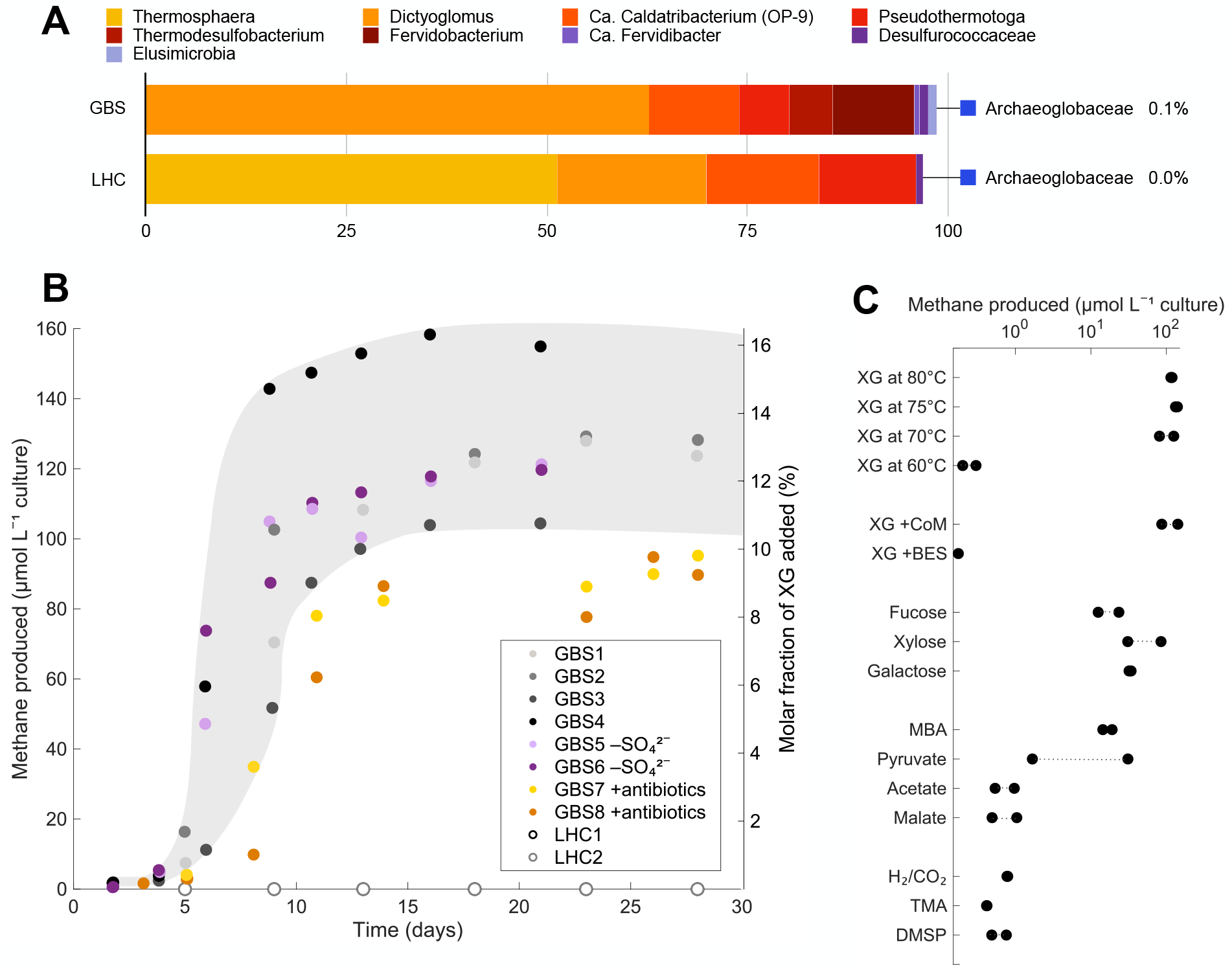
Community composition and methane production data of the xyloglucan-degrading culture.**A** Community composition (%) in anaerobic enrichments from Great Boiling Spring (GBS) and Little Hot Creek (LHC) based on 16S rRNA gene tags. Taxa are identified at the lowest named rank according to Silva. All *Archaeoglobaceae* sequences detected belong to *M. nevadense*. **B** Headspace methane concentration over time. The second y-axis shows the relative amount of methane derived from the organic carbon added. The antibiotics mix consisted of streptomycin and carbenicillin to specifically target bacteria. Approximate range in values from four GBS cultures is colored in gray. SO_4_^2−^, sulfate. **C** Headspace methane concentrations in duplicate treatments on day 15. If not specifically indicated, incubation temperature was maintained at 75 °C. Methoxybenzoate, MBA; Trimethylamine, TMA; Dimethylsulfoniopropionate, DMSP.

After transfers into fresh media, we henceforth measured methane production in microcosms and assessed the influence of potential stimulants and inhibitors. The XG enrichment from LHC did not produce methane at any time during the experiment, while the GBS culture reached 104-158 µmol methane L^−1^ after two weeks of incubation at 75 °C, which appeared to be the optimal temperature for methane production (Fig. 1B, C). At that time, there were 1.2-2.0 × 10^8^ cells mL^− 1^ culture based on direct total cell counts. Methane formation at 80 °C was not significantly lower than that at 75 °C (Student’s t-test, P < 0.05) and we also observed methane in cultures grown at 85 °C (data not shown). Methanogenesis in the *Archaeoglobi* may, hence, increase the known temperature limit of terrestrial methanogenesis by 5-10 °C (Hedlund *et al*., 2015). We henceforth refer to the culture from GBS (and not LHC) as XG-degrading culture. The omission of sulfate as a potential oxidant for anaerobic respiration and the addition of antibiotics targeting bacteria resulted in only minor differences in methane formation (Fig. 1B). Amendment with coenzyme M (CoM), an essential nutrient for some methanogens (Taylor *et al*., 1974) and one species of non-methanogenic *Archaeoglobus* (Mori *et al*., 2008), did not increase activity or growth. In contrast, addition of bromoethanesulfonate (BES), a structural analog to methyl-CoM and commonly used methanogenesis inhibitor specifically targeting Mcr, halted methane production completely (Fig. 1C). We then tested if monosaccharide products of XG breakdown, specifically fucose, xylose, or galactose, would stimulate methane production in the absence of XG, however, less than half the amount of methane was produced (with the exception of one xylose replicate reaching ∼85 µmol methane L^−1^). Interestingly, incubations with methoxybenzoate (MBA) supported 14-19 µmol methane L^−1^, suggesting the capacity for methoxydotrophic methanogenesis (Mayumi *et al*., 2016). Other organic acids and well-known methanogenic substrates (H_2_/CO_2_, TMA, DMSP) resulted in low methane formation (< 10 µmol methane L^−1^), with ambiguous results from pyruvate incubations (Fig. 1C).

### A functional, divergent Mcr in *M. nevadense*

To better understand the pathway of methane generation in *M. nevadense*, we carried out metagenome sequencing of our enrichment cultures. Despite the low abundance of *M. nevadense* as determined by 16S rRNA gene sequencing, we were able to recover a MAG estimated to be 63 % complete with 2.4 % contamination (*M. nevadense* XG-1). GTDB_tk placed this MAG within the genus “WYZ-LMO2” (Wang *et al*., 2019), members of which were reported to contain Mcr operons and originally named *Candidatus* Methanoproducendum (Hua et al., 2019) or later described with the synonym *Candidatus* Methanomixophus (Liu *et al*., 2020). Phylogenomic analysis of currently available *Archaeoglobi* genomes placed our MAG together with “Bin 20”, or *M. nevadense* GBS^Ts^, which was derived from an *in-situ* AFEX-treated corn stover enrichment (IMG: 3300005298_ 20) (Peacock *et al*., 2013). These two MAGs represent the same species, sharing 98.4 % average nucleotide identity (Yoon *et al*., 2017) with a tetra-nucleotide index of 0.996 (Table S1). Since the 1.6 Mb-comprising MAG from the *in-situ* enrichment was far more complete (98 % and 3.7 % contamination), the genomic analysis below focused on *M. nevadense* GBS^Ts^. Screening the unassembled metagenomic reads for the *mcrA* gene using GraftM revealed a gradual enrichment of *M. nevadense* from 0.3 % in GBS sediment, over 10 % in the *in-situ* cellulolytic enrichment (corn stover bag), to 100 % in the XG enrichment culture (Fig. 2A), suggesting that while a diversity of Mcr-containing organisms is present *in-situ, M. nevadense* was the sole source of methane in our incubations. The phylogenetic placement of the recruited McrA is close to the “WYZ-LMO2” McrA on a monophyletic branch with protein sequences derived from Korarchaeota, Nezhaarchaeota, and Verstraetearchaeota (Wang *et al*., 2019) (Fig. 2B). Although the McrA shows affiliation with Archaea outside of the Euryarchaeota, a genome tree derived from an alignment of 43 genomic markers placed *M. nevadense* among other Euryarchaeota within the family *Archaeoglobaceae* with strong support. Based on monophyly and ANI values > 95 % (Jain *et al*., 2018), MAGs derived from GBS belong to a single species in the genus *Methanoproducendum* with *M. nevadense* GBS^Ts^ as the nomenclatural type for the species (Fig. 2C, Table S2), whereas genomes representing other species in the genus have been recovered from Yellowstone National Park or Jinze Hot Spring in Tengchong, China (Hua *et al*., 2019; Wang *et al*., 2019; Liu *et al*., 2020). To promote best practices in systematics, a protologue for this new species is included in the Supplements, and *M. nevadense* together with *Methanoproducendum hydrogenotrophicum* (type is Bin16^Ts^), *Methanoproducendum dualitatem* (type is LMO3^Ts^), and *Methanoproducendum nevadense* (type is GBS^Ts^) will be registered as species belonging to the genus *Methanoproducendum* in the SeqCode Registry (Hedlund *et al*., 2022).

**Fig. 2.**
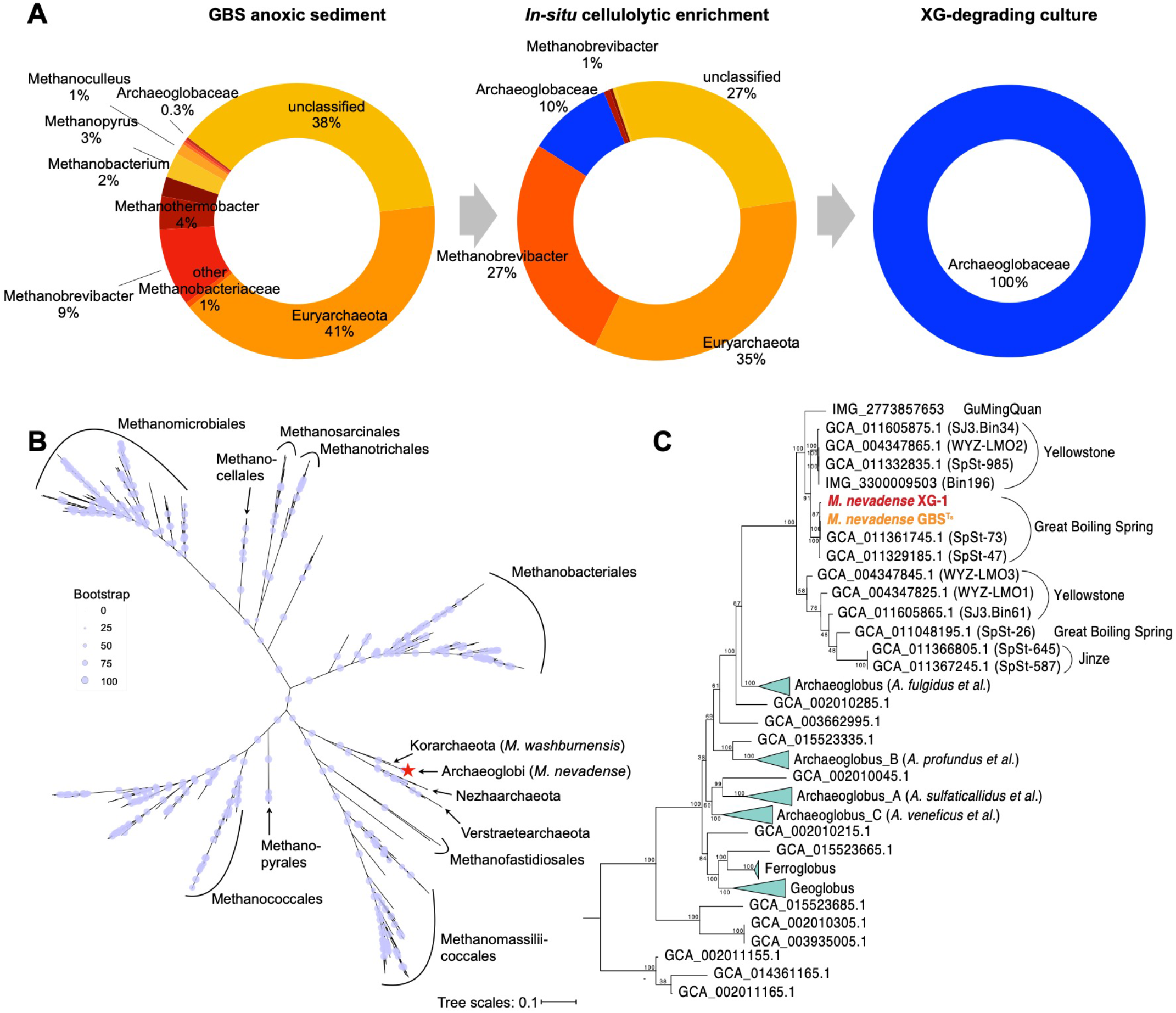
Enrichment of *M. nevadense* among multiple McrA-encoding taxa in GBS. **A** Genus-level classification of *mcrA* gene sequences in the original GBS sediment, the *in-situ* cellulolytic enrichment, and culture, following an enrichment trend of *M. nevadense*. **B** Phylogenetic relationship between *M. nevadense* McrA (red star) and 499 related McrA amino acid sequences aligned by PSI-Blast. **C** Genome tree of the two *M. nevadense* MAGs and 78 other *Archaeoglobi* genomes. *M. nevadense* forms a distinct branch with McrA from other species of *Methanoproducendum*.

### Potential carbon and electron transfers in *M. nevadense* methanogenesis

The *M. nevadense* MAG encodes several methane-specific metabolic features that could support a variety of pathways for methane generation and energy conservation. Most conspicuous are the Mtr and Mcr operons, which are only known to function in methane-metabolizing organisms, catalyzing the penultimate and final steps of methanogenesis (Fig. 3). Besides these operons, the Wood-Ljungdahl (WL) pathway is largely the same as in other *Archaeoglobi* and could be used in either the anabolic direction for acetyl-CoA generation or in the catabolic direction for multicarbon substrate breakdown. An important difference is absence of the main subunit of Mer (Methylenetetratetrahydromethanopterin reductase), which carries out one of the key C1 oxidation/reduction steps in this pathway (Fig. 3). This absence was reported in a previous analysis of similar MAGs (Liu *et al*., 2020), but we add here that two homologs of Mer are encoded in this genome (as well as cultured *Archaeoglobus* strains). Mer and its homologs are part of the large luciferase-like monooxygenase family (pfam00296) and it is possible one of these paralogs is able to complete the C1 pathway between CO_2_ and methyl oxidation states. However, an incomplete WL pathway could explain the inability of *M. nevadense* to use H_2_ as a sole methanogenic substrate in our experiments.

**Fig. 3.**
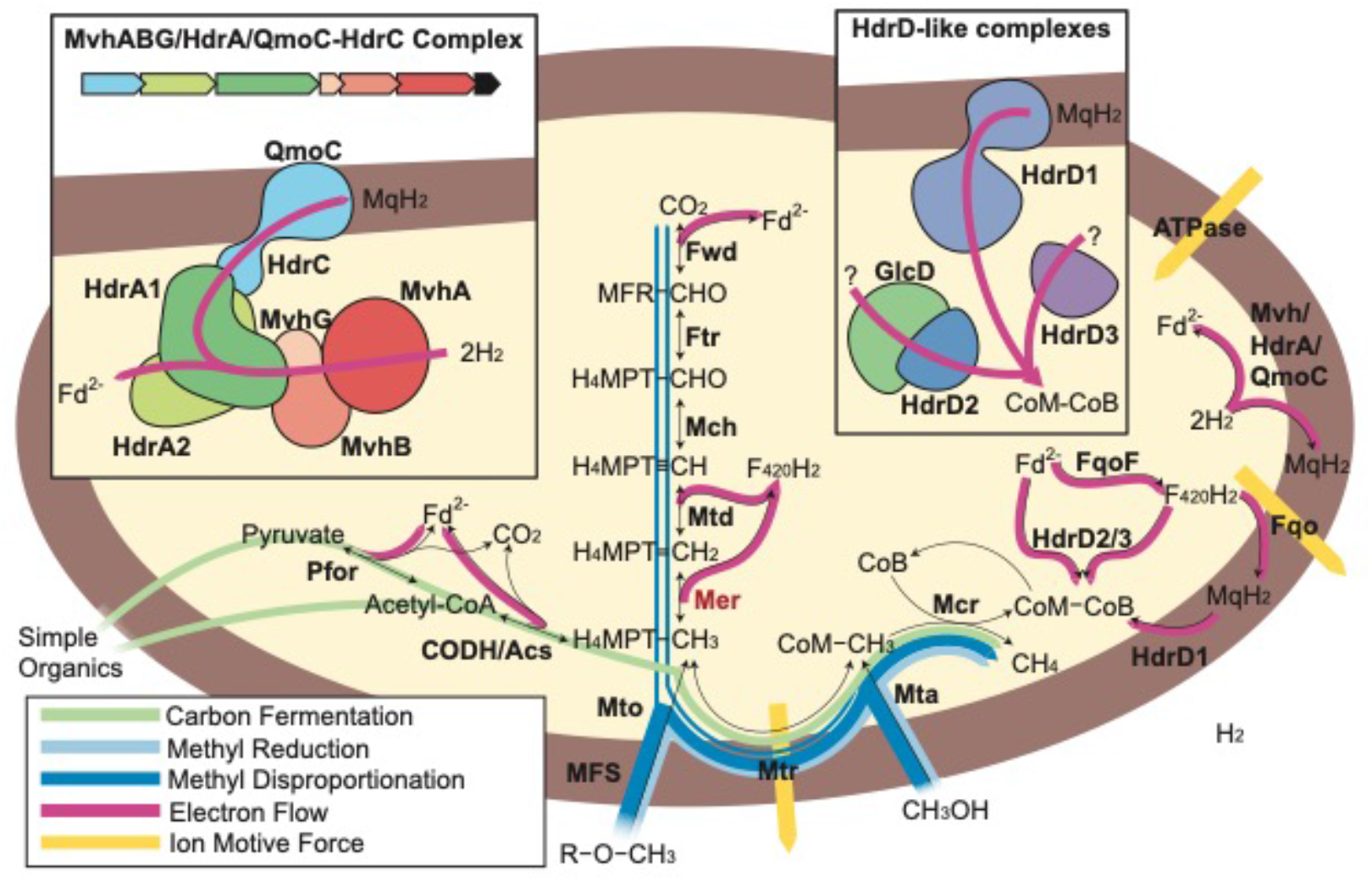
Possible methanogenesis pathways in *M. nevadense*. Key genes encoded in the *M. nevadense* GBS^Ts^ MAG are shown along with pathways for carbon and electron flow that would be consistent with our substrate incubation experiments. All genes present in the genome are shown in bolded black letters, including six of the seven traditional methanogenesis pathways (Fwd, Ftr, Mch, Mtd, Mtr and Mcr, see Table S3). The ability to incorporate methyl groups from methanol and methoxylated compounds is enabled by the Mta and Mto complexes, respectively. Methyl groups could be used for methanogenesis on their own via methyl disproportionation (dark blue), or methyl reduction with electrons from hydrogen (light blue). An additional possibility is the degradation of more complex carbon compounds for carbon and electrons (light green). Electron flow (pink) to the CoM/CoB heterodisulfide could be supported by three HdrD homologs found in the MAG (inset, top right). The HdrD2-GlcD gene cluster is found together with the Mtr operon and associated methanogenesis genes, making this the most likely candidate. Another conspicuous electron transport gene cluster is highlighted in the top right, consisting of a NiFe hydrogenase, two HdrA homologs, and a membrane-bound cytochrome b HdrC-QmoC fusion expected to interact with the menaquinone pool. The flow of methyl groups through Mtr (depending on the direction) and electron flow through Fqo generates an ion motive force (yellow), ultimately generating ATP via ATP synthase.

Methyl groups on H_4_MPT or CoM could be reduced to methane in two ways. First, six electrons could be generated by oxidizing a methyl group to CO_2_ through the WL pathway, which could be used for the reduction of three other methyl groups to methane (Methyl disproportionation, dark blue, Fig. 3). This would rely on the distant homologs of Mer as described above. Second, electrons sourced from hydrogen could be used to reduce methyl groups to methane (Methyl reduction, light blue, Fig. 3). A third possible route of carbon into this pathway could be through the breakdown of multicarbon compounds via pyruvate and acetyl-CoA. These carbons would end up as CO_2_ (in oxidation reactions producing reduced Fd) and methyl groups on H_4_MPT (green path, Fig. 3).

There are a variety of electron transfer complexes encoded in the *M. nevadense* MAG that could support electron flow from Fd, F_420_H_2_ or H_2_ onto HdrD. An Fqo complex very similar to the one characterized in *A. fulgidus* is found in the genome and would be able to transfer F_420_H_2_ electrons onto menaquinone (Brüggemann *et al*., 2000). Fd electrons are the lowest potential, and could possibly pass to F_420_H_2_ through soluble FqoF or directly onto Fqo *sans* Fqof, both of which have been proposed as options in cultured methanogens (Welte and Deppenmeier, 2011). A fascinating gene cluster containing the [NiFe] hydrogenase subunits MvhABG, two HdrA homologs, and a fusion of QmoC and HdrC is found in the *M. nevadense* MAG and is absent in the cultivated non-methanogenic *Archaeoglobi*. The QmoABC complex is normally found in *Archaeoglobi* and other sulfate reducers and is thought to carry out electron transfer from menaquinone to AprAB during sulfate reduction (Duarte *et al*., 2016). AprAB is absent from *M. nevadense* along with the other key sulfate reduction proteins Sat, DsrAB and DsrMKJOP. QmoABC has been proposed to bifurcate electrons (Appel *et al*., 2021), and in the absence of its recognized partners in QmoAB or AprAB, it is tempting to speculate this complex is involved in electron flow between H_2_, ferredoxin and menaquinone.

These electron flow reactions would likely be associated with the generation of a proton motive force, either through vectoral proton pumping at Fqo, or through quinol loops at the b-type cytochromes in the QmoC homologs. Also, depending on the direction of carbon flow through Mtr, additional sodium motive force maybe be generated at this step. Ion motive force ultimately would be used for the generation of ATP through ATP synthase.

### XG-degrading culture consumes H_2_ concomitant to methane production

Because of the apparent discrepancy between the genetic potential to consume H_2_ for methanogenesis and the very low activity observed in H_2_/CO_2_ incubations, we tested if the enriched microbial community produces H_2_ and if there would be a difference in H_2_ with the application of BES. If Mcr-based methanogenesis consumed H_2_, there would be more H_2_ available in BES-inhibited cultures.

Indeed, 12 to 20 days after XG incubation we detected H_2_ in the culture headspace, indicating H_2_ production by members of the culture. We furthermore observed more than twice the H_2_ accumulation when BES was added (Fig. 4). This implies H_2_ consumption during methanogenesis by the XG-degrading culture and suggests *M. nevadense* is co-dependent on H_2_ and on another, unknown compound (see discussion).

**Fig. 4.**
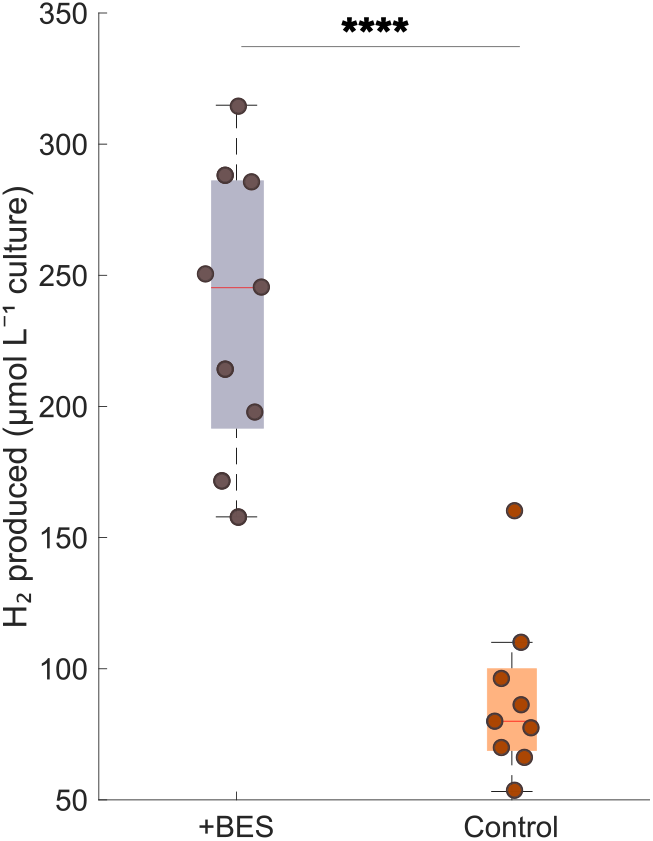
Headspace H_2_ concentration in triplicate cultures between day 12 and 20. H_2_ was significantly more abundant in +BES incubations (ANOVA, \*\*\*\**p* < 0.0001, n = 9). Central marks in boxes indicate the median, while the bottom and top edges are the 25^th^ and 75^th^ percentiles. Whiskers extend to the most extreme data points not considered outliers.

### A microbial origin of GBS methane

To evaluate the influence of biological methane, possibly from *M. nevadense*, on the overall methane pool dissolved in the hot spring water, we collected dissolved methane from seven hydrothermal pools in the Great Boiling Spring geothermal field (Fig. S1) and determined the C and H isotope composition. These values reveal a primarily volcanic/sedimentary thermogenic origin of the dissolved methane from all pools (Fig. 5A), given that lower values of δ^13^C and δ^2^H are associated with biogenic processes (Schoell, 1988). In contrast to microbial methanogenesis, thermogenic methane production is an abiotic process, although thermogenic methane originates from biologically produced material, i.e., high molecular weight carbon (Schoell, 1988; Etiope *et al*., 2011). However, the δ^13^C of methane decreases by up to ∼10 ‰ in individual pools of 48-94 °C (GBSTb, GBSTa, G04b, GBS19), suggesting the potential contribution of hydrogenotrophic methane production at these sites (Bradley and Summons, 2010). The more positive δ^2^H values in methane from spring G04c perhaps result from mixing with methane from the air due to less upwelling water movement (>1.5 days residence time in G04 springs, (Costa *et al*., 2009)). We complemented the isotope data with methane formation rates from *in-situ* anoxic sediment incubations in a subset of pools. Biogenic methane would be consistent with a significant methane accumulation over time.

**Fig. 5.**
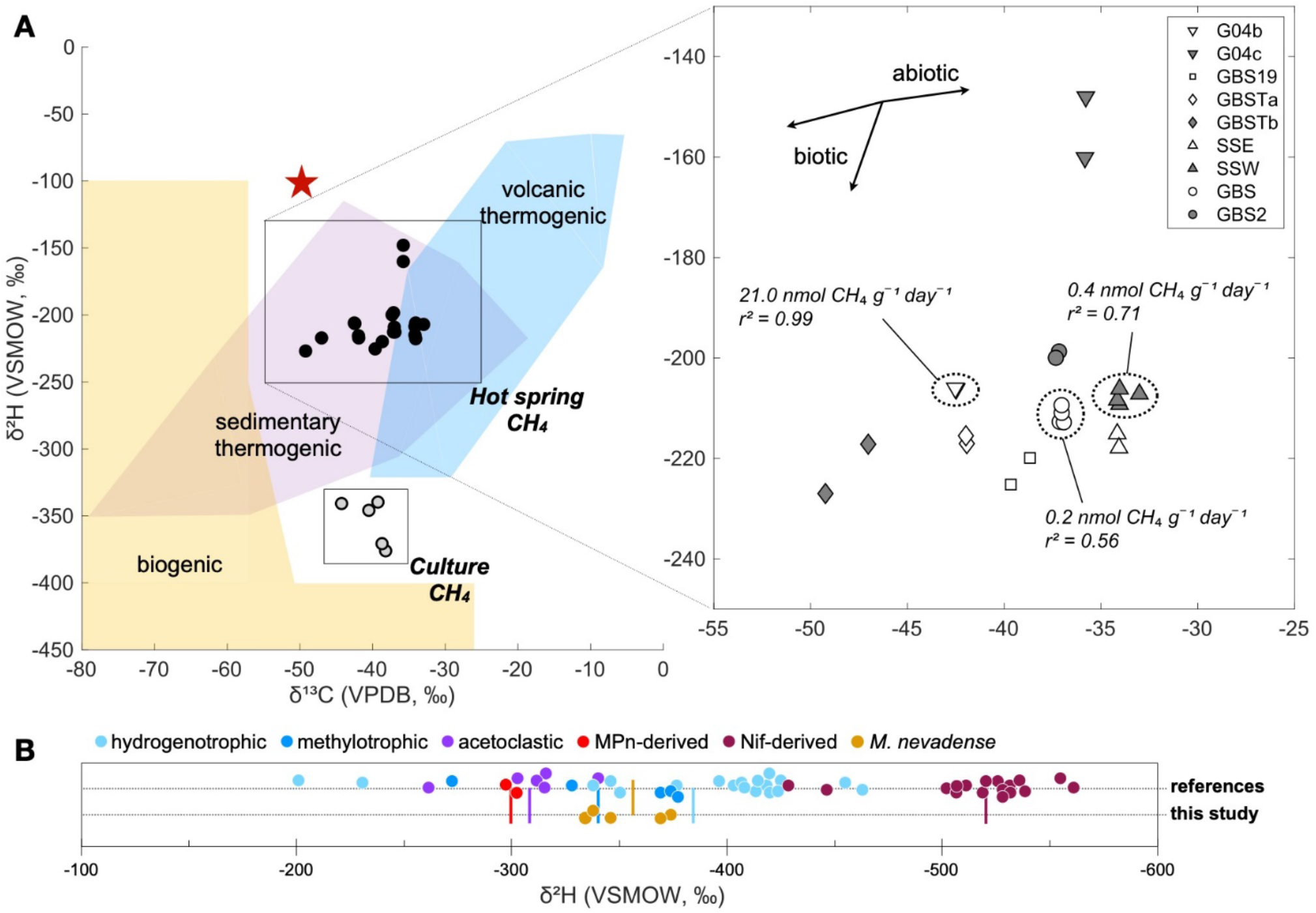
Isotopic composition of *in-situ* and culture methane gas in a δ^13^C-*δ* ^2^H space. **A** Known ranges of values (Schoell, 1988) may indicate the origin of methane at GBS. Atmospheric methane is indicated by a red star as reference. Rates in the inset were derived by hot spring *in-situ* incubations. See Fig. S2 for location information and basic geochemical data on individual hot springs. **B** *δ* ^2^H of methane from culture headspace and reference data from hydrogenotrophic, methylotrophic, acetoclastic (Gruen *et al*., 2018), methylphosphonate (MPn)-derived (Taenzer *et al*., 2020), and nitrogenase (Nif)-derived (Luxem *et al*., 2020) methane production pathways. Colored vertical lines indicate means in *δ* ^2^H of individual pathways.

Reflecting the more negative δ^13^C results at G04b, methane production here was > 20 nmol methane g^−1^ sediment day^−1^ higher than in SSW or GBS (Fig. 5A, inset). Although this supports the idea that biological methane influences total dissolved methane abundances through mixing, a higher sample size would be needed to corroborate this trend. Methane produced by the XG-degrading culture was indistinguishable from GBS methane in its ^13^C composition but exhibited a strong depletion in ^2^H, conforming with its biological origin. Because of this difference, we compared δ^2^H values in methane from various biogenic pathways revealing a large data spread across studies (Fig. 5B). Regardless of this variability, methane putatively produced by *M. nevadense* resembled most closely that originating from methylotrophic and hydrogenotrophic pathways based on δ^2^H. Overall, our isotopic analysis shows a dominance of abiotic methane in GBS waters, with potential contributions from methyl- and H_2_-dependent microbial methane production at some sites.

## Discussion

We present evidence that a previously uncultured member of the *Archaeoglobaceae, M. nevadense*, harbors a divergent Mcr and is capable of methane production (Figs 1 and 2). Our calculated cell-specific methanogenesis rates of 78.9-176 fmol cell^−1^ day^−1^ reside between 0.5 fmol cell^−1^ day^−1^ (Beulig *et al*., 2019) and 576-43,200 fmol cell^−1^ day^−1^ (Ver Eecke *et al*., 2013; Topçuoğlu *et al*., 2016; 2019; Stewart *et al*., 2019) derived from pure cultures of mesophilic (lower boundary) and hyperthermophilic (upper boundary) deep-sea methanogens.

Methanogenic rates mediated by *Archaeoglobi* are therefore competitive with rates mediated by other Euryarchaeota. Based on the cultivation experiments, we postulate four possible modes of methane formation in the XG-degrading culture (Fig. 6), two of which we can rule out as explanations for our observations.

**Fig. 6.**
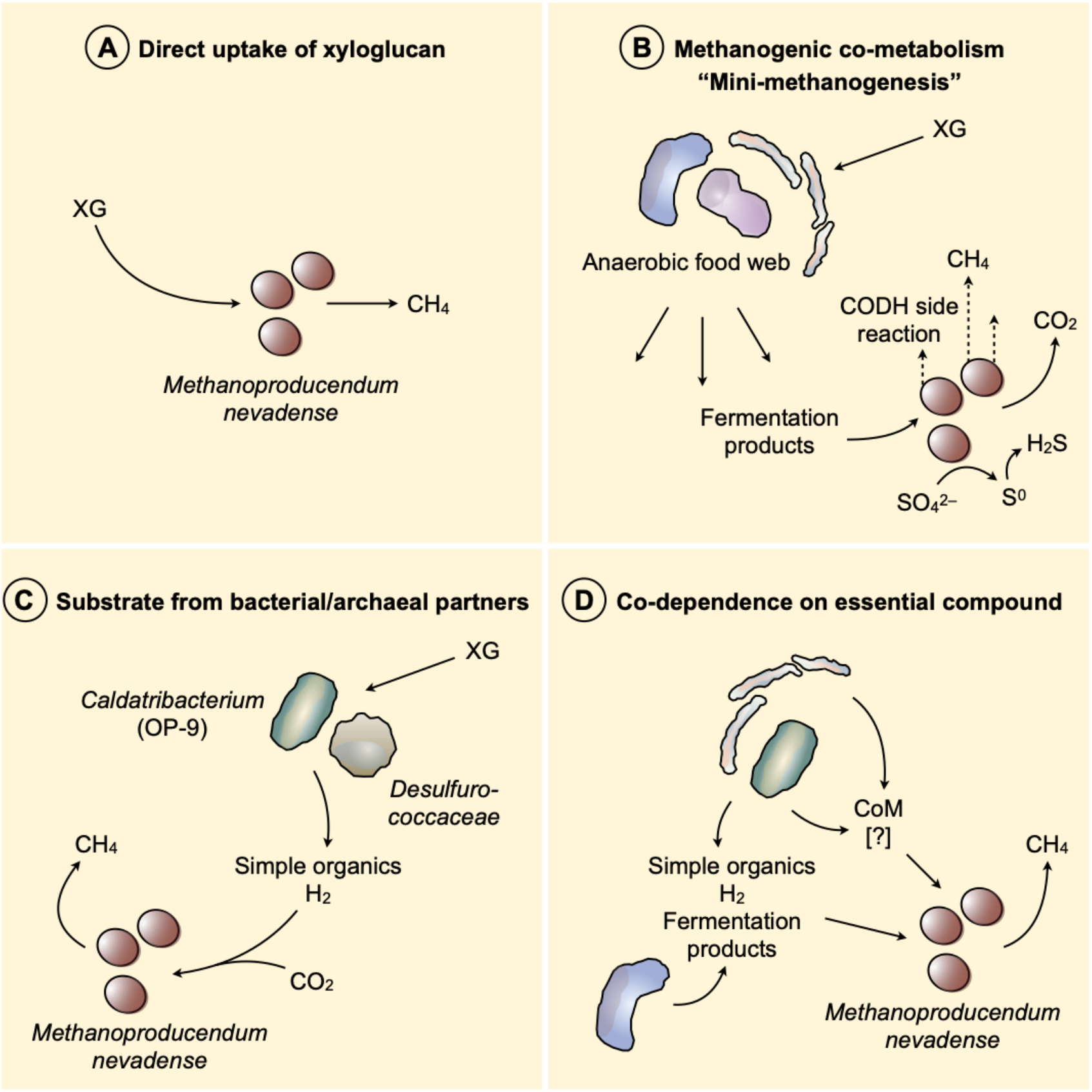
Schematic illustration of postulated modes of methane formation by *M. nevadense* in the xyloglucan enrichment culture. Fermenters other than depicted here are possible (see text). Mechanisms in panels C and D are considered more likely than those in A and B. CODH, carbon monoxide dehydrogenase.

First, *M. nevadense* directly depolymerizes XG to mineralize the resulting sugars into methane gas (Fig. 6, panel A). Methane accumulation in the presence of bacterial antibiotics and favorability of XG over all other substrates tested would suggest this mode, however, the *M. nevadense* genome does not indicate any genes typically associated with XG breakdown, including glycoside hydrolases (Kampik *et al*., 2021), rendering direct XG metabolism unlikely.

Second, the observed methane is produced as an accidental byproduct of other biochemical processes, also described as “mini-methanogenesis” (Fig. 6, panel B). Several isolated *Archaeoglobus* strains are known to produce methane in an Mcr-independent process. Sulfate-reducing strains of *Archaeoglobus fulgidus* are capable of making methane at < 200 µmol L^−1^ culture (Stetter *et al*., 1987; Beeder *et al*., 1994) and the sulfite-reducing *Archaeoglobus veneficus* produces methane at 0.6-2 µmol L^−1^ culture (Huber *et al*., 1997). *Archaeoglobus infectus*, isolated from an active submarine volcano in the Western Pacific, also reduces sulfite and generates methane at ∼5 µmol L^−1^ culture (Mori *et al*., 2008). The source of the trace amounts of methane in these pure cultures is thought to be a spurious side-reaction of carbon monoxide (CO) dehydrogenase (Vorholt *et al*., 1995; Klenk *et al*., 1997). Methane production in these *Archaeoglobus* cultures may serve to export excess reducing equivalents rather than for conserving energy. *M. nevadense*’s complete Mcr gene cluster (Fig. 3) and the observation that methane production is blocked by BES addition (Fig. 1C) robustly suggests that Mcr-independent mini-methanogenesis is not responsible for methane production in *M. nevadense*.

Third, *M. nevadense* lives in a syntrophic relationship with partner bacteria or archaea (Fig. 6, panel C). *Ca*. Caldatribacterium, *Ca*. Fervidibacter, *Fervidobacterium, Dictyoglomus*, or *Desulfurococcaceae*, each genomically predicted or known to depolymerize polysaccharides, could degrade XG to simple organics (XG monomers) and H_2_ that is recycled to methane by *M. nevadense*. This mode would make XG degradation more energetically favorable under conditions with limited terminal electron acceptors. The lower but sustained methane production in the presence of antibiotics argues against strict reliance on bacterial partners and suggests an archaeal-archaeal partnership with yet-uncultivated *Desulfurococcaceae* (Graham *et al*., 2011). The fact that the XG monomers fucose, xylose and galactose stimulated methane production (Fig. 1C), albeit not to the level of XG, is also supportive of that proposed mode. Cultured members of the *Archaeoglobaceae* metabolize a much broader range of carbon substrates than cultured methanogens. There is no methanogen known that can grow by fermenting pyruvate or glucose, for example. Our genome analysis (Fig. 3) highlights multiple potential pathways for energy conservation from methanogenesis in *M. nevadense* by using a variety of methylated compounds on their own or with electrons sourced from H_2_ or the fermentation of more complex carbon compounds. Comparing the amount of methane produced (Fig. 1A) to the amount of H_2_ consumed (Fig. 4) by the enrichment culture, we note that the molar H_2_/methane ratio we measure (∼1.2) deviates from the theoretical stoichiometric ratio expected if methanogenesis is strictly hydrogenotrophic (4). It is possible that methanogenic *Archaeoglobi* occupy a more metabolically versatile niche as compared to traditional methanogens, where they can utilize a wide range of carbon substrates generated from complex carbon breakdown supplemented with H_2_ consumption.

Fourth, *M. nevadense* metabolizes fermentation products derived from XG degradation but is also dependent on other compounds provided by the microbial community, such as vitamins and cofactors (Fig. 6, panel D). This mode would explain our observation that culture-derived H_2_ was consumed by methanogenesis, whereas exogenous H_2_ in the absence of XG or other organic carbon was not (Fig. 4 versus Fig. 1C). As a refinement of the third mode, the methanogen could still live in syntrophy with another archaeal species. CoM may be such a cofactor obligately required for methane metabolism (Gunsalus and Wolfe, 1977; Mori *et al*., 2008), together with vitamins such as folate (Buchenau and Thauer, 2004) or amino acids. However, in our experiments, CoM addition to XG-grown cultures did not stimulate methanogenesis, suggesting additional dependence on an essential compound and/or an already saturated supply of CoM by other members of the community.

Concerning the question about the origin of methane at GBS, our data suggest that it is primarily thermogenic (abiotic), in line with methane fluxes correlating with temperature in GBS pools (Hedlund *et al*., 2011). The range in δ^13^C of our measurements (Fig. 5A) is congruent with data collected in pyrolysis experiments at 400 and 500 °C involving montmorillonite clays (Sackett, 1978). As reaction mechanism, carbon bond cleavage via an acid-activated carbonium ion at the clay mineral surface has been proposed (Greensfelder *et al*., 1949). Such a mineral-catalyzed mechanism would be feasible within the sedimentary deposits at GBS that are composed of clays from the Eastern Sierra Nevada (Birkeland and Janda, 1971). The location of the culture methane in δ^13^C-δ^2^H space extends the coverage of biogenic methane, and we propose that methane retrieved from thermal systems exhibiting similar δ^13^C-δ^2^H values to be considered biogenic in origin. Comparing our δ^2^H data to reference data derived from methane produced by well-characterized thermophilic methanogens (Fig. 5B) supports the use of methyl groups and H_2_ inferred from our genomic analysis.

## Conclusion

We conclude that *M. nevadense* is a methanogen with a highly versatile and possibly syntrophic lifestyle. In addition to using sugars and organic acids to fuel methanogenesis, our experimental (methane production with MBA addition) and genomic data (a complete MtoABCD complex) suggest *M. nevadense* is capable of methoxydotrophic methanogenesis (Mayumi *et al*., 2016), a methanogenic pathway not previously demonstrated in thermophiles. This metabolic flexibility enables *M. nevadense* to cooperate with diverse microbial partners and to adapt to a wide array of organic carbon sources. Because of the close relationship between the *M. nevadense* Mcr complex and others from diverse uncultivated archaea (Fig. 2B), we propose that the similarly divergent Mcr complexes in Korarchaeota, Nezhaarchaeota, Verstraetearchaeota, and other *Archaeoglobi* are also functional and likely generate methane (Vanwonterghem *et al*., 2016; Berghuis *et al*., 2019). Our findings may also expand methane metabolism to a wider range of hot spring chemistries (Reigstad *et al*., 2010). Taken together, our study represents a first step to experimentally verify the genomic potential contemplated for a variety of Mcr-encoding archaea. Future studies will further unravel the significance of such lineages in the evolution of methane metabolism and as source of biological methane on Earth.

## Data Availability

Metagenome and 16S rRNA gene sequence data will be deposited in the NCBI database upon publication.

## Acknowledgements

We thank Marike Palmer for advice on systematics, Alma Parada for discussions about the computational approach, Alexander Jaffe for feedback on the manuscript, David and Sandy Jamieson for access to Great Boiling Spring, Alexander Manriquez for help with sampling at GBS, and Michael Morikone for initial metagenome analysis of the xyloglucan culture. We appreciate the CSUSB Genomics course of Spring 2018 for assistance with xyloglucan culture DNA extraction and long read sequencing.

## Author Contributions

SB and AED developed the experimental design with input from BPH and JAD. SB conducted the experiments with assistance from MEQ. JAD achieved enrichment of *M. nevadense* and has maintained the XG-degrading culture long-term. Metagenomic work was done by SB and GLC. GLC performed the metabolic modeling. SB and JAD conducted field work at GBS. SB wrote the paper with contributions from GLC, BPH and AED, and all authors edited the final version.

