## Supplementary Text and Figures for "Mcr-dependent methanogenesis in *Archaeoglobaceae* enriched from a terrestrial hot spring"

*for*

SI material includes:

| | |
|---|---|
| Supplementary Text | Page 2 |
| Protologue | Page 3 |
| Figure S1 | Page 4 |
| Figure S2 | Pages 5 & 6 |
| Table S1 | Supplementary data file 1 |
| Table S2 | Supplementary data file 2 |
| Table S3 | Supplementary data file 3 |

**Supplementary Text: Carbon and electron transfer pathways in *M. nevadense* GBS$^{Ts}$**

Two methyltransferase systems are present in the *M. nevadense* GBS$^{Ts}$ MAG that are likely inputs for C1 moieties to the methanogenesis pathway. The first, MtoABCD, was recently characterized in detail in *Archaeoglobus fulgidus* and enabled growth on methoxylated compounds such as 2-methoxyphenol. The second is a homolog of the MtaCAB methyltransferase system specific for methanol. These methyltransferases will bring in C1 moieties at the same oxidation state, but importantly for the bioenergetics of the pathway, on opposing sides of the Mtr complex. Mto, which is present in *A. fulgidus* that lacks CoM, transfers methyl groups onto $H_4MPT$. These methyl groups, if used for methanogenesis, would pass through the Mtr complex to CoM, which would conserve energy in the form of a sodium motive force. The Mta complex transfers methyl groups directly to CoM, bypassing this energy conserving step.

Once methane is generated by Mcr, electrons must flow to CoM-CoB to complete the pathway. These electrons will be found on a mix of Fd, $F_{420}H_2$, or $H_2$, depending on which of the above carbon transformations are carried out. The heterodisulfide reductase reaction takes place in an active site found in HdrD/HdrB homologs in normal methanogens. Only HdrD-like versions of these proteins are found in *M. nevadense* and are found alone (HdrD3), fused to membrane-integral cytochrome B domains (HdrD1), or in association with other, poorly characterized electron transfer proteins (HdrD2). HdrD1 and 3 are found in cultured, non-methanogenic *Archaeoglobi*, while HdrD2 is found in a large gene cluster with many methanogenesis-specific proteins such as Mtr and McrA2, and is not found in cultured *Archaeoglobi*, suggesting this might be the main source of electrons for CoM-CoB reduction.

**Protologue**

Description of *Methanoproducendum nevadense* sp. nov.

*Methanoproducendum nevadense* [ne.va.den'se] L. masc./fem. adj. *nevadense*, from the US state of Nevada

Anaerobic, thermophilic (≤ 70°C) archaea that inhabit circumneutral, terrestrial geothermal systems and produce methane by using a methyl-CoM reductase complex. Hydrogen is used as an electron donor. Methoxybenzoate is likely used as a methyl donor for methanogenesis. Products of sugar fermentation are likely methanogenic substrates. Genomes belonging to this species are ~1.6 Mb and have a single copy of rRNA genes. Genomic assemblies for this species originated from sediments from Great Boiling Spring in Nevada, USA. The nomenclatural type for the species is the genome GBS$^{Ts}$ (Need accession number).

**Supplementary Figures**

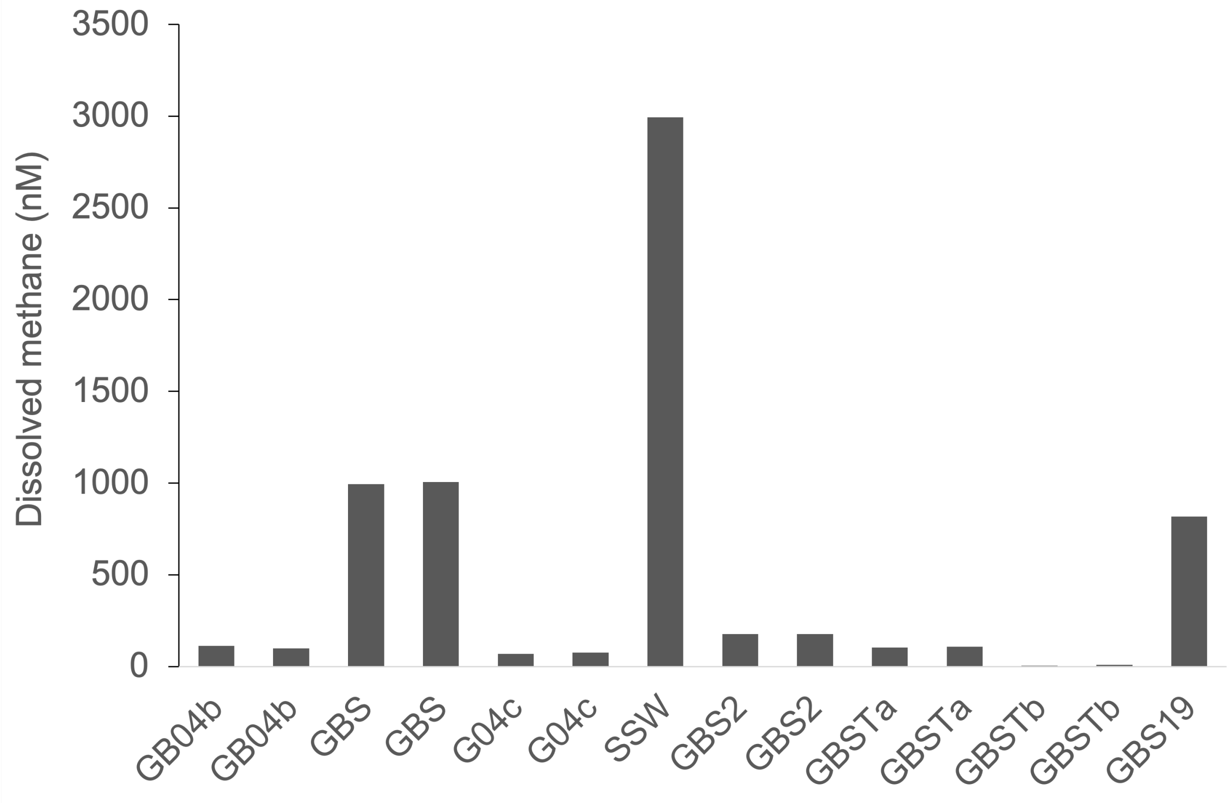


**Figure S1.** Dissolved methane concentrations in hot spring pools of the GBS geothermal field. Some measurements were conducted on replicate (unique) water samples. All samples were retrieved from 5-20 cm water depth.

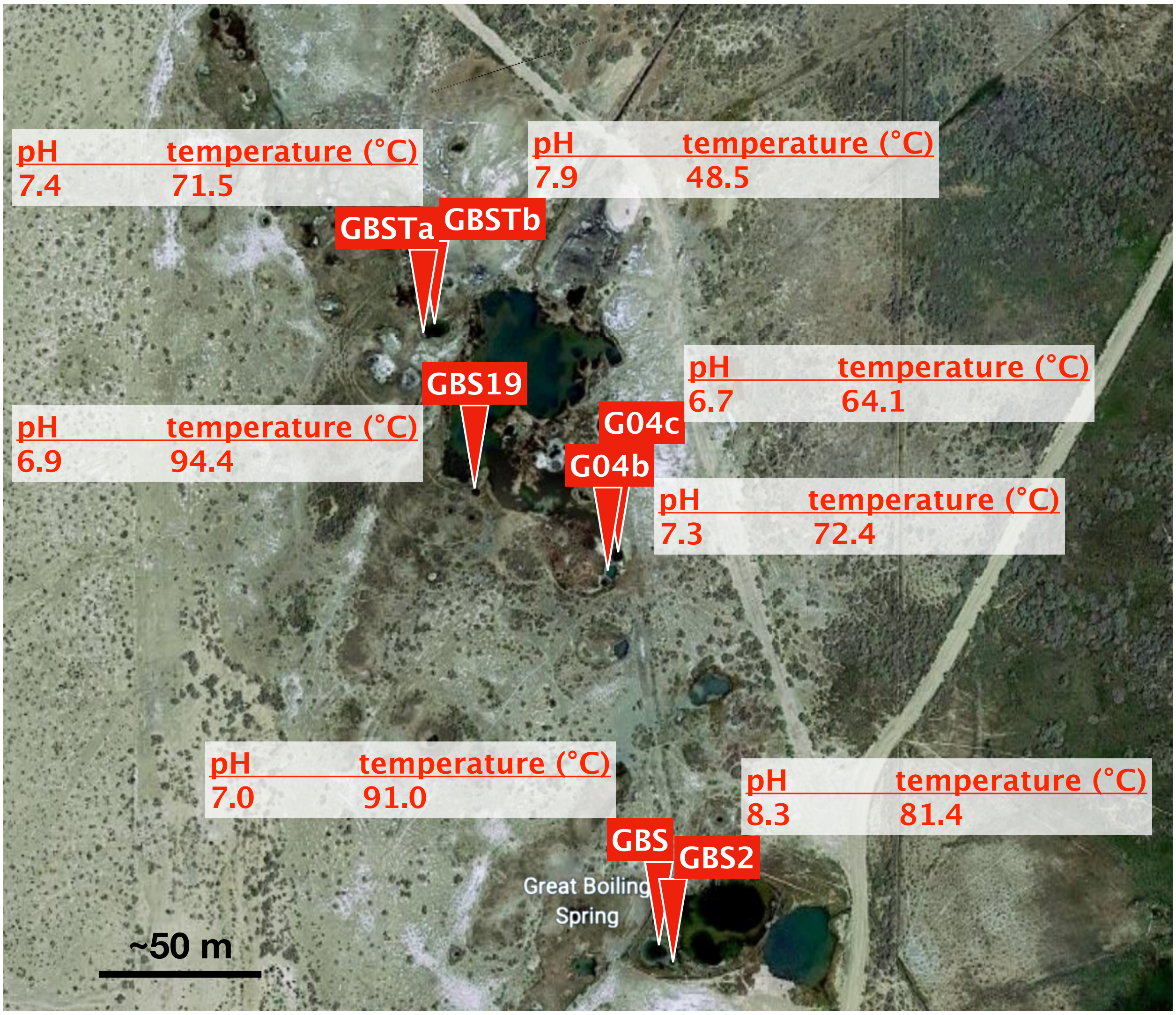


**Figure S2.** Location and pH/temperature of GBS hot spring pools studied. Top of the map points north. GBS is the Great Boiling Spring main pool, with GBS2 being a small outflow from the main pool. The G04 pools are located adjacent to each other and are connected by surface streams. GBS19 is a deep and very hot pool that emits much water steam. GBST is a low temperature pool with vegetation on its shore (*Juncus*-dominated). Sampling point GBSTa was in the middle of the pool and GBSTb was close to the shore above green microbial mats. Map retrieved from Apple Maps version 2.1.

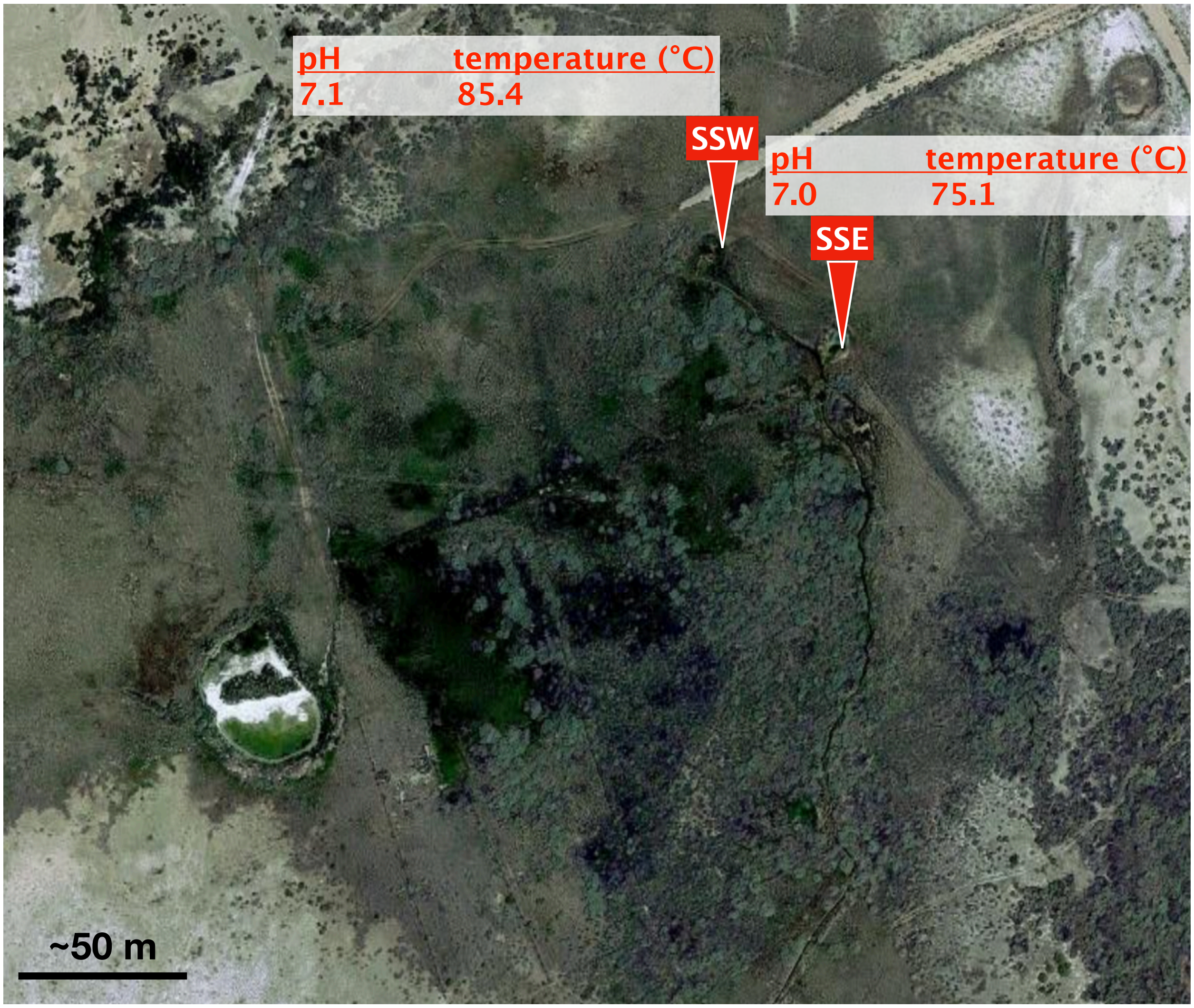


95 **Figure S2 cont.** Location and pH/temperature of GBS hot spring pools studied. Top of the map points north. SSW (Sandy Spring West) is a small pool enclosed by vegetation and is feeding into SSE (Sandy Spring East) by a ~40 m creek. The Sandy Springs are located ~600 m south-west of the GBS main pool. Map retrieved from Apple Maps version 2.1.
